# Large-scale clinical interpretation of genetic variants using evolutionary data and deep learning

**DOI:** 10.1101/2020.12.21.423785

**Authors:** Jonathan Frazer, Pascal Notin, Mafalda Dias, Aidan Gomez, Kelly Brock, Yarin Gal, Debora S. Marks

**Affiliations:** Department of Systems Biology, Harvard Medical School, Boston, MA 02115, USA.; OATML Group, Department of Computer Science, University of Oxford, Oxford, OX1 3QD, UK.; Broad Institute of Harvard and MIT, Cambridge, MA 02142, USA.

## Abstract

Quantifying the pathogenicity of protein variants in human disease-related genes would have a profound impact on clinical decisions, yet the overwhelming majority (over 98%) of these variants still have unknown consequences^1–3^. In principle, computational methods could support the large-scale interpretation of genetic variants. However, prior methods^4–7^ have relied on training machine learning models on available clinical labels. Since these labels are sparse, biased, and of variable quality, the resulting models have been considered insufficiently reliable^8^. By contrast, our approach leverages deep generative models to predict the clinical significance of protein variants without relying on labels. The natural distribution of protein sequences we observe across organisms is the result of billions of evolutionary experiments^9,10^. By modeling that distribution, we implicitly capture constraints on the protein sequences that maintain fitness. Our model EVE (Evolutionary model of Variant Effect) not only outperforms computational approaches that rely on labelled data, but also performs on par, if not better than, high-throughput assays which are increasingly used as strong evidence for variant classification^11–23^. After thorough validation on clinical labels, we predict the pathogenicity of 11 million variants across 1,081 disease genes, and assign high-confidence reclassification for 72k Variants of Unknown Significance^8^. Our work suggests that models of evolutionary information can provide a strong source of independent evidence for variant interpretation and that the approach will be widely useful in research and clinical settings.

## Introduction

The exponential growth in human genome sequencing has underlined the substantial genetic variation in the human population. Understanding the clinical relevance of this genetic variation has the potential to transform healthcare and motivates the massive government investments in the collection of human population genomic information together with demographics and clinical data such as the UK BioBank^1^, ChinaMAP^24^, deCODE^25^. This access to sequencing has enabled both genetic studies that associate variants with diseases and more mechanism-based approaches that associate variants with biochemical and cellular phenotypes. However, relating specific changes in the genome to disease phenotypes remains an open challenge as the number of variants in the human population dwarfs the number we are able to investigate. Protein coding regions alone contain large variation between people; 6.5 million missense variants have been observed to date (gnomAD^2^) and the consequences of the vast majority (98%) of these, even in disease related genes, are unknown^3^. It is estimated that there will be a variant for every protein position (bar lethal) somewhere in the 9 billion human population.

Given this challenge, new high-throughput experimental technologies have emerged that are able to assess the effects of thousands of mutations in parallel (sometimes called Deep Mutational Scans (DMSs) or Multiplexed Assays of Variant Effects MAVEs)^11–23,26,10,27^. However, these technologies do not easily scale to the whole human proteome, especially not to combinations of variants, and critically depend on assays that are relevant and mappable to human diseases.

Ideally, computation could accelerate clinical variant interpretation. However, widely-used computational methods^4–7^ are *supervised*, training on sparse, imbalanced and noisy clinical labels^28^, and can be implicitly circular due to data leakage during cross-validation^29^ (Methods). This leads to questions about the advertised accuracy that may not hold for out-of-training variants and genes, hence compromising generalizability and scope^29^. It is hardly surprising, therefore, that there is some hesitance to use computational methods for anything but “weak evidence”^3^, as in the guidelines for variant classification from The American College of Medical Genetics and Association for Molecular Pathology (ACMG-AMP)^8^.

By contrast, *unsupervised* probabilistic models of evolutionary sequences alone have been remarkably successful at predicting the effects of variants on protein function and stability^30,31 9 32–34^ and are fundamentally generalizable as they avoid learning from labels. However, whilst they have achieved state-of-the-art performance for predictions across hundreds of thousands of mutation effect experiments^10,35^, they have not been developed to address the issue of clinical relevance. To do this for all human proteins, methods must be scalable, robust across protein families and be able to map evolutionary fitness to a pathophysiological score.

We introduce EVE (Evolutionary models of Variant Effects), a new computational method for the clinical classification of human genetic variants trained solely on evolutionary sequences. We show via careful validation that EVE predicts the clinical significance of variant effects, outperforming current state-of-the-art computational predictors (without the risk of overfitting clinical labels), and is surprisingly as accurate as many high-throughput experiments. We make predictions for 11 million variants in 1,081 disease associated genes and subsequently assign high-confidence reclassification of over 94k of Variants of Unknown Significance.

## Results

### Unsupervised models accurately predict clinical significance labels

Our method – EVE – is designed to learn from sequence variation across species the propensity of human missense variants to be pathogenic, and it does so in two main steps (Fig. 1). First, to successfully capture the constraints from evolution as observed in extant sequences, which manifest as complex dependencies across positions, and learn a distribution over the corresponding sequence space, we need a sufficiently expressive generative model. To that end, we leverage Variational Autoencoders (VAE), which have been successful in learning complex high-dimensional distributions across multiple domains^36,37^ including predicting protein function^10^. For each human protein of interest, a Bayesian VAE is trained on a multiple sequence alignment retrieved by searching ~250 million natural protein sequences from *O*(10^5^) organisms in UniRef^38^ (Methods, Supplementary Table 1). We optimized model architecture and training hyperparameters via thorough ablation analyses, and demonstrate its superiority over prior methods (Methods, Extended Data Fig. 1 and Extended Data Fig. 2). By sampling from the approximate posterior distribution of the VAE, we obtain the “evolutionary index” for each single amino acid variant, which estimates the relative likelihood of the variant with respect to the wild type (Methods). We compared this evolutionary index against clinical labels and observed that the score that separates pathogenic from benign labels is remarkably consistent across proteins (Fig. 2a), suggesting we may use unsupervised clustering methods to predict the propensity of a variant to be pathogenic entirely without learning from previous clinical information. Therefore, in the second step, rather than using supervised learning to map scores to label categories, we fit a two-component global-local mixture of Gaussian Mixture Models^39^ on the distributions of evolutionary indices for all single amino acid variants across proteins (Methods, Extended Data Fig. 3). The output of this process is both the EVE score – a continuous pathogenicity score defined over the interval [0,1], with zero being most benign and one being most pathogenic – and class assignments. For these assignments, after computing the predictive entropy to identify variants for which the predictions are most uncertain, we bin variants into one of three categories: Benign, Uncertain, Pathogenic (Methods).

**Figure 1.**
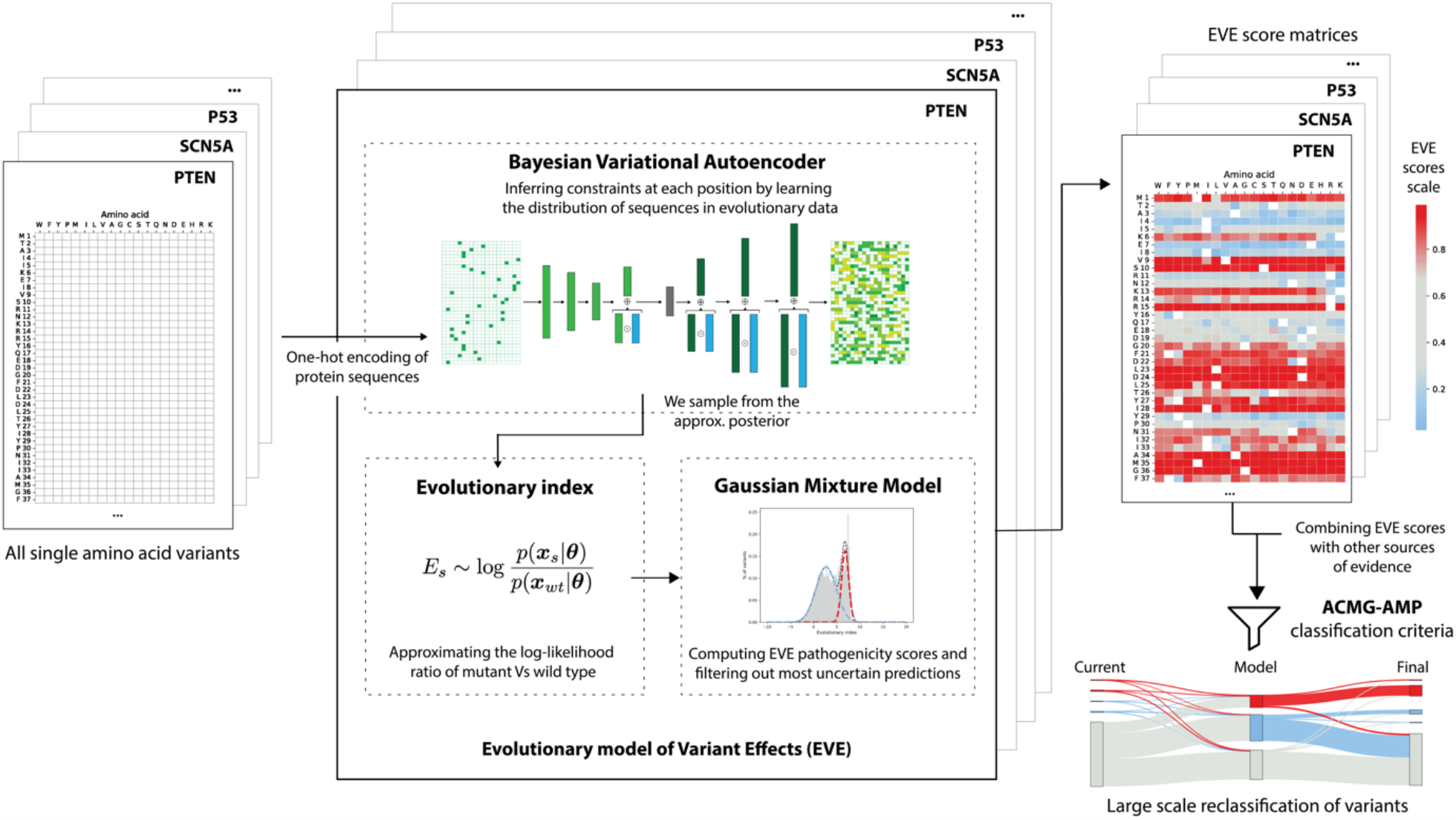
EVE Overview. *EVE* is a set of protein-specific models providing for any single mutation of interest a score reflecting its propensity to be pathogenic. For each protein, a Bayesian VAE learns a distribution over amino acid sequences from evolutionary data. It enables the computation of the *evolutionary index* for each mutant, which approximates the log-likelihood ratio of the mutant vs the wild type. A global-local mixture of Gaussian Mixture Models separates variants into benign and pathogenic clusters based on that index. The EVE scores reflect probabilistic assignments to the pathogenic cluster. When combining EVE scores with other sources of evidence following ACMG-AMP^8^ guidelines, and taking EVE as a “strong source” of evidence, we can classify over 4x more benign and pathogenic variants than are currently known in ClinVar^3^.

**Figure 2.**
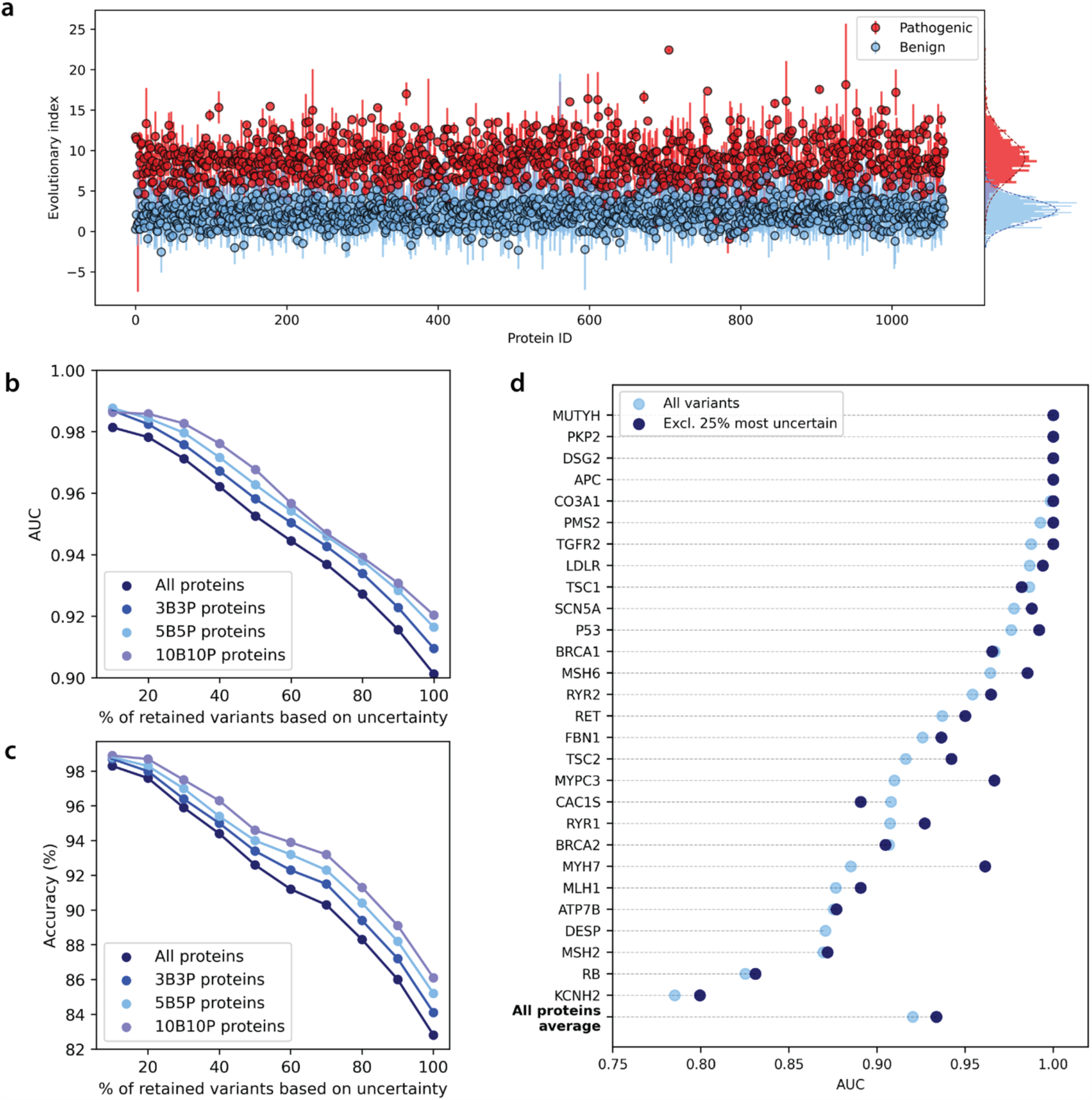
EVE performance. **a**. Average evolutionary indices per protein, and corresponding standard deviations, for variants with known Benign and Pathogenic clinical labels across 1,081 proteins (sorted by alphabetical order). On the right, marginal distributions over the 1,081 proteins. **b**. AUC per protein of the EVE score against clinical labels, for the subset of ACMG “actionable genes”^40^ with at least 5 Benign and 5 Pathogenic labels. AUCs are computed both for EVE scores of all variants, and of the 75% variants with most confident scores (Methods). The last row shows the AUC computed over all variants and over the 75% with most confident scores, for 1,081 proteins. **c. & d**. AUC and accuracy of our model, respectively, as a function of the percentage of variants used for calculation, based on the uncertainty of their EVE score. Both AUC and accuracy are computed for 1,081 proteins, and for the subsets thereof with 3, 5 and 10 Benign and 3, 5 and 10 Pathogenic labels – denoted as 3B3P, 5B5P and 10B10P, respectively.

We apply EVE to a set of human genes that have been associated to human disease (1,081), collectively containing 11 million variants (Methods). To validate our model, we compute the area under the receiver operating characteristic curve (AUC) and accuracy with respect to clinical labels of the whole 1,081 proteins, both in aggregate and per protein, as well as test sensitivity to the number of labels per protein (Fig. 2b, 2c, Supplementary Table 2). EVE pathogenicity scores are predictive of clinical labels, with an aggregate AUC of 0.90 and mean AUC per protein of 0.92. The performance is robust to the number of labels per protein (Fig. 2b and 2c) demonstrating generalizability to genes and variants with less or no annotation, as we would expect from an unsupervised approach. To show specific examples we selected the subset of 28 genes from the ACMG “actionable genes” list^40^ with at least 5 Benign and 5 Pathogenic labels (Fig. 2d). Of these, nine have an AUC ≥ 0.99 (APC, MUTYH, DSG2, PKP2, CO3A1, PMS2, TGFR2, LDLR, TSC1).

EVE outperforms state-of-the-art supervised methods (MISTIC^6^, M-CAP^5^, Polyphen2^4^). A key reason for using a fully unsupervised approach is to bypass the biasing issues and difficulties in cross-validation due to data-leakage faced by supervised methods (Methods) ^29^. These issues can lead to an overestimation of the supervised performance when using metrics such as AUC and accuracy. Nonetheless, EVE achieves higher performance across proteins based on these two metrics (Extended Data Fig. 4, Supplementary Table 3).

The EVE score reflects the probabilistic assignment to either pathogenic or benign clusters. The probabilistic nature of the model enables us to quantify our uncertainty on this cluster assignment, which implies we can bin variants into Benign and Pathogenic categories as conservatively as we wish by assigning some variants as Uncertain. As uncertainty correlates with modelling mistakes, increasing the percentage of variants assigned Uncertain leads to increased AUC and accuracy (Fig. 2b, 2c, Supplementary Table 2). By setting the 25% of variants with highest classification uncertainty as Uncertain, we attain an overall AUC of 0.93 (mean AUC per protein 0.93) and 90% accuracy. Our method enables practitioners or researchers to adjust this proportion of excluded variants and select specific trade-offs between accuracy and coverage for different disease scenarios. To aid these decisions, and since AUCs can mask asymmetries in the relation between precision and recall (Extended Data Fig. 5), beside summary performance metrics we also provide detailed performance curves (ROC and precision-recall) for each gene (see data availability).

Having validated our model’s performance at predicting clinical significance, we can now compare its performance to that of high-throughput experimental assays, which are themselves increasingly being used as a source of evidence for variant classification.

### Model as predictive of clinical significance as high-throughput experiments

New technologies (DMS and MAVEs) that measure the effect of 10^3^-10^4^ variants simultaneously are being adopted to accelerate our understanding of human genetic variants^11–23^. A direct comparison of our model predictions to that of MAVEs for 17 assays on 7 genes^11–13,17,20,23,41–45^ indicates very similar performance in predicting clinical significance as measured by AUC. Using a variety of label definitions (Methods, Supplementary Table 4), we find EVE to outperform MAVEs on average (Extended Data Fig. 6, Supplementary Table 5).

For the five genes with MAVE assays that also have high quality ClinVar labels – BRCA1, P53, PTEN, MSH2, SCN5A – the overall performance of EVE at predicting clinical significance is again as good as, or better than, that of experiments (Fig. 3, Extended Data Fig. 7, Methods). Impressive technology for saturation genome editing of BRCA1 by Findlay et al. was specifically motivated by the need to classify newly observed variants as more of the population is sequenced, and had good agreement with previous ClinVar missense labels, demonstrating the utility of these high-throughput experiments for novel classifications^28^. Both Findlay et al and EVE achieve an AUC of 0.95 against 52 labels of the RING and BRCT domains of BRCA1. Similarly, Jia et al^41^ motivated their deep mutational scan of MSH2 (a DNA mismatch repairs gene associated with Lynch syndrome) by the need to classify thousands of variants which have uncertain significance. Here, our model predicts clinical significance better than the experimental data^41^ (AUC 0.86, 0.82 respectively over 93 labels). The same is true for the tumor protein P53 (AUCs 1 (EVE), 0.9 (experiment^20^) over 97 labels) and for the tumor suppressor PTEN (AUCs 0.99 (EVE), 0.99 (experiment^13^) over 106 labels and 0.97 (EVE) and 0.88 (experiment^43^) over 66 labels). In the case of the Brugada syndrome ion channel gene SCN5A the number of labels, 8, is too small to compute reliable AUCs, but these labels are perfectly separated by both EVE scores and experimental fitness metrics^15^ (Supplementary Table 5, Methods).

**Figure 3.**
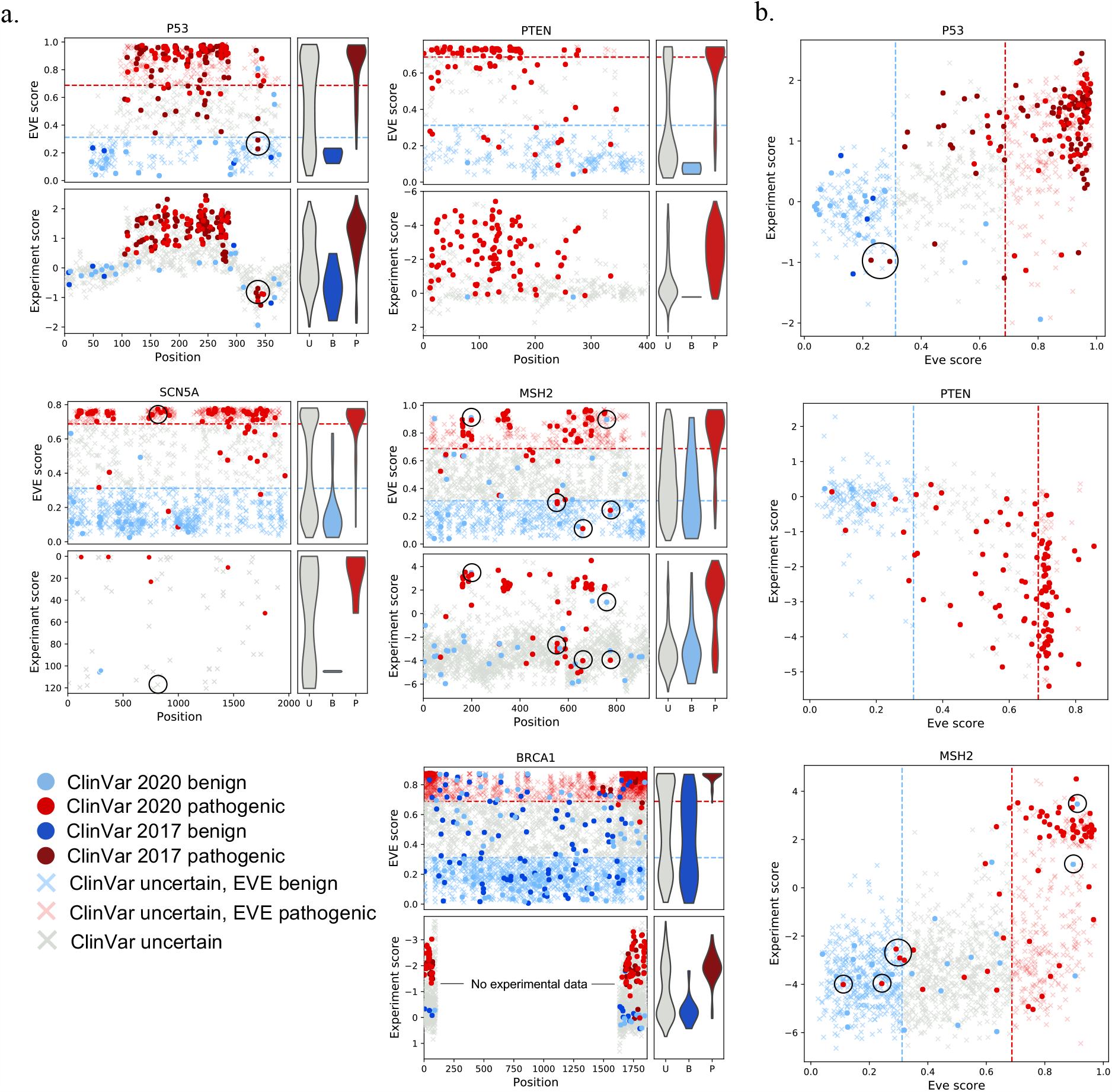
Computational model EVE as good as high-throughput experiments for clinical labels. **a**. Comparison of computational model predictions (upper panels, y-axis is EVE score) and experimental assay measurements (lower panels, y-axis experimental assay metric) to ClinVar labels (dots) and VUS (crosses), where pale red and pale blue crosses indicate EVE assignments of previous VUS mutations. Dashed red and blue lines correspond to the 75% confidence threshold of EVE scores. x-axes are position in protein. Violin plots show scaled distributions of VUS, Benign and Pathogenic labels. Experimental measurements data from deep mutational scans of P53^20^, PTEN^13^, MSH2^41^, BRCA1^12^and patch clamp of ion channel for SCN5A ^14^. Circled are R337H/C in P53, S554N/T, D660G, I774V, E198G, G759E in MSH2 and R814Q in SCN5A. **b**. Scatter plots comparing experiment scores to EVE score.

A more in-depth analysis further reveals situations where model and experiments have distinct strengths and weaknesses. For instance, Giacomelli et al P53 assay^20^ achieves perfect separation of benign and pathogenic variants up until near the “tetramer domain” (C-terminus) whereas EVE predictions achieve near perfect separation throughout (Fig. 3a upper left). Another example is R814Q in SCN5A which Glazer et al. found to have near wild-type peak density in their patch clamp assay and yet has been observed in multiple cases of both Brugada syndrome and long QT syndrome. They hypothesize this to be a gain of function phenotype^15^. EVE indeed predicts this variant to be pathogenic, supporting their hypothesis (Fig. 3a middle left). This suggests there may be a more general distinction between MAVEs and our model – while MAVEs can probe either gain or loss of specific functions, EVE scores are based on evolutionary constraints and hence do not distinguish mechanisms, instead measuring the variant’s effect on the fitness of the organism.

Interestingly, since EVE and MAVEs are independent sources of evidence, comparison of their results may help evaluate the clinical labels themselves. 85% of the 27 variants where EVE score disagrees with ClinVar across MSH2, PTEN and P53 are supported by MAVE experimental data (Fig. 3). For example, both EVE and experimental assays support a benign score where ClinVar has Pathogenic labels for variants R337H and R337C in P53, S554N/T, D660G and I774V in MSH2, and 15 variants in PTEN. Similarly, both EVE and experimental assays support a pathogenic clinical effect where ClinVar has Benign labels for G759E and E198G in MSH2 (the pathogenic assignment of the latter is further supported by new experimental data^46^).

Taken together, our analysis shows that our computational method EVE can predict clinical significance as well as high-throughput experiments, but it also highlights how experiments and model, being independent sources of evidence, can work synergistically towards an understanding of the genetic basis of human disease.

### Clinical interpretation of 11 million variants

The primary hurdle to using computational methods to interpret variants at scale is validation. In the previous sections, through careful benchmarking, we have made the case that deep unsupervised models of evolutionary data provide strong evidence for assessing the clinical significance of variants. Before performing classification (see below), we provide the continuous EVE scores for all single variants for all 1,080 proteins together with browsable mutation matrices, allowing for more nuanced evaluation (Fig. 4a, see data availability).

**Figure 4.**
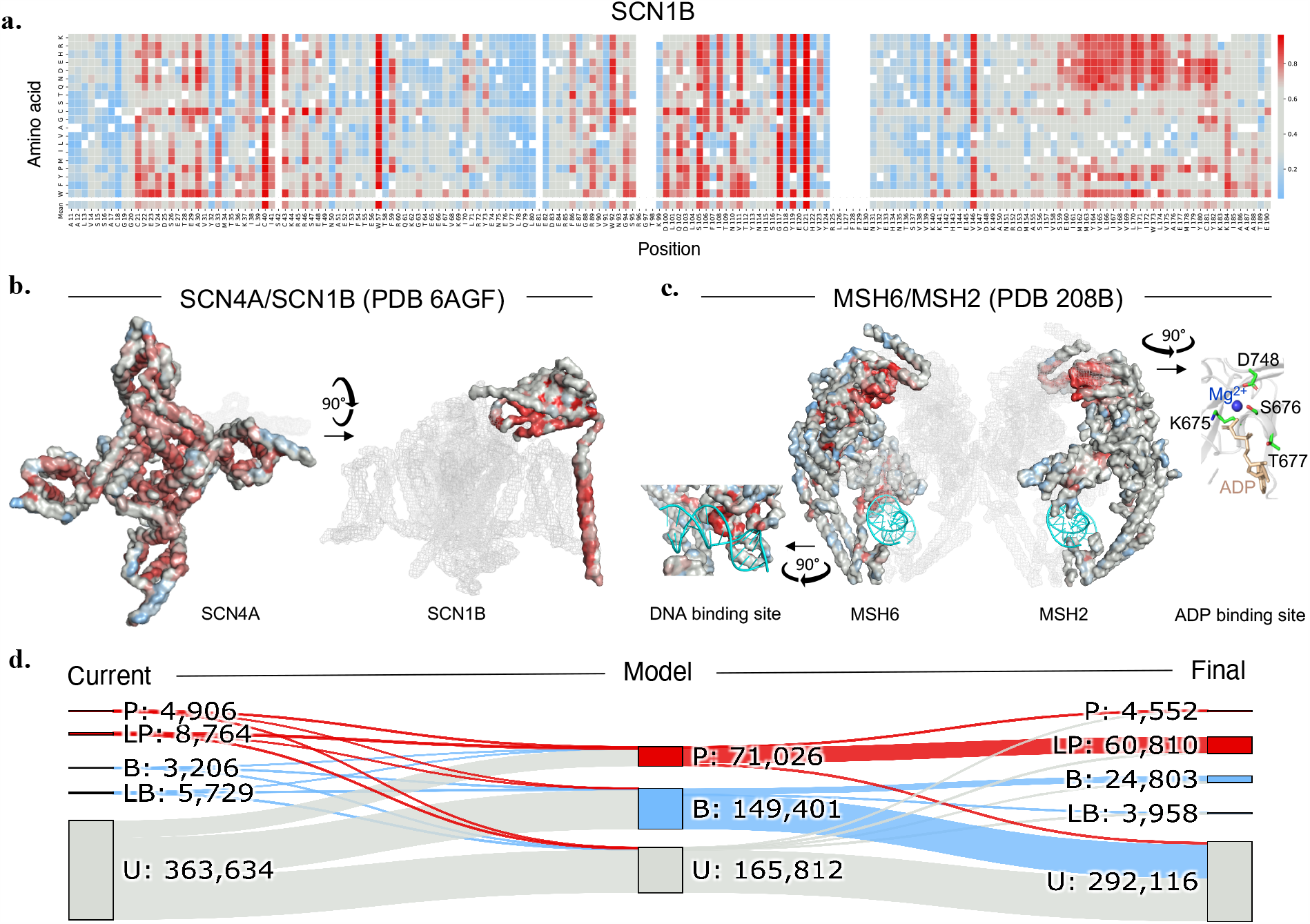
Clinical interpretation of 386,239 variants in 1081 genes. **a**. Heatmap of EVE pathogenicity scores in SCN1B. **b**. and **c**. Representations of 3D structures of SCN4A/SCN1B (PDB 6agf ^56^) and MSH6/MSH2 bound to ADP and DNA (PDB 2o8b^57^), colored by mean score per position (SCN4A, MSH6, MSH2) and maximum score per position to show bimodal scores in TM-helix (SCN1B). Clusters of high pathogenicity scores in 3D include the pore region of SCN4A, the TM-helix and the hydrophobic core residues in the IgG domain of SCN1B; the ADP binding site of MSH2 (D748N/V/H, K675E, S676L, T677R (VUSs) and the DNA binding region of MSH6. **d**. Evolutionary model (EVE) enables ACMG-AMP-like classification of variants at scale. The total number of (Likely) Pathogenic labels (13,670), with at least a one-star rating, in ClinVar for the 1,081 proteins (left, red). The model classifies 71,026 variants as pathogenic (middle, red), which when combined with other sources of evidence in an ACMG-AMP-like manner, results in 65,362 variants which we recommend classifying as (Likely) Pathogenic (4.8x increase) (right, red). The equivalent analysis for (Likely) Benign labels yields 8,935 in ClinVar (left, blue), 149,401 predicted by model (middle, blue) and 28,761 which we recommend classifying as (Likely) Benign (3.2x increase) (right, blue).

Many of these proteins have strong links with disease but only a fraction of the variants seen in the population have known associations. For instance, variants in genes SCN4A, SCN5A and SCN1B in the ion channel complex have been associated with neurological, cardio-vascular and muscular disorders ^47–50^ but less than 10% of the ~3,000 variants seen in the population have a clinical interpretation. Similarly, variants in MSH2 and MSH6 of the mismatch DNA repair complex are associated with hereditary nonpolyposis colorectal cancer^51^ and with ~20% of sporadic cancers^52^, but only 4% of observed variants have clinical labels (Extended Data Table 1).

The distribution of EVE scores across the proteins and the amino acids in each position follows trends that might be expected by the relative functional importance of that region in the protein, such as clusters of regions in 2D along the linear sequence that coincide with secondary structure^9^, the hydrophobic cores, ligand-binding and active sites. For example, in the SCN4A/SCN1B complex, many variants at the interface between the two proteins, variants lining the SCN4A pore and specific positions in the hydrophobic core of IgG domain of SCN1B are predicted strongly pathogenic with high confidence (Fig. 4a, 4b). Existing clinical labels are much sparser and often only single variants in a position were known (e.g. C121W and V138I pathogenic and benign respectively in SCN1B). Similarly, amongst the strongest EVE pathogenic signals for MSH2 are variants proximal to the bound ADP and Mg^2+^ in the known 3D structure (D748N/V/H, K675E, S676L or T677R). In MSH6, beside the corresponding ADP binding site, variants at residues proximal to the bound DNA have strong pathogenicity scores (Fig. 4c).

In addition to providing continuous scores, our model is well suited for performing classification. A significant advantage of the model being trained on evolutionary data alone (as compared to models which train on multiple sources of evidence^4,5,29^) is that the EVE score may be regarded as providing evidence from a single source. It is therefore straightforward to incorporate it into a variant classification scheme such as the one proposed by ACMG-AMP, which seek to establish a systematic procedure for combining orthogonal sources of evidence. The 1,081 disease related genes in our analysis have 387k variants which have been observed in at least one human to date.

Of these, about 94% are variants of uncertain significance and only the remaining 6% have been classified as any of Benign, Likely Benign, Likely Pathogenic, Pathogenic (Fig. 4d, left). For the purpose of classification, the EVE scores may be grouped into three categories – Benign, Pathogenic and Uncertain, after accounting for modelling uncertainty (Methods). Here we report results with 40% of variants set as Uncertain, which corresponds to a 90% overall accuracy performance. Of the 60% assigned clinical significance, ~70% are Benign and ~30% are Pathogenic (Fig. 4d, middle, Methods).

The final classifications of our analysis (Fig. 4d, right) results from combining model classifications with population data from gnomAD and other forms of evidence, weighted according to the ACMG-AMP guidelines, while treating the EVE scores as providing “strong” evidence (Methods). This is the same strength as a well-established functional assay, which seems reasonable given the comparisons to MAVEs we have presented. The result is that 17% of variants seen in the human population are classified as (Likely) Pathogenic, 7% as (Likely) Benign and 76% as Uncertain, representing a four-fold increase in the number of known (Likely) Benign or (Likely) Pathogenic missense variants. The vast majority of the Pathogenic assignments by EVE get classified as (Likely) Pathogenic (92%), in contrast to just 19% of the EVE Benign assignments getting a final classification as (Likely) Benign. This is a reflection of the ACMG-AMP criteria, and highlights the asymmetry between pathogenic and benign assignments – it is easier to establish with genetics that a variant is involved in disease than to establish the opposite. This asymmetry, however, is not present in a model based on evolutionary constraints like ours (Fig. 4a, Supplementary Table 6).

### Systematic approach to flag clinical labels that may be incorrect

There are 2,192 variants for which model assignments contradict the current ClinVar status, and we found supporting evidence for EVE’s predictions for 164 of them (Fig. 4, Supplementary Table 7). This highlights a further benefit of our approach – that by being trained on independent evidence and being unsupervised, our model provides a natural means of detecting potentially incorrect labels (Methods). No (Likely) Pathogenic ClinVar labels with one or more review status stars were in this subset of 164, demonstrating excellent agreement between our analysis and high-quality (Likely) Pathogenic labels. Our algorithm did however flag as possibly incorrect 121 of ClinVar (Likely) Benign labels thought to be of high quality. Examples include MSH2 variants described earlier in this text and P53 variant R337Q (Fig. 3).

## Discussion

We have demonstrated that deep unsupervised models trained on sequence alignments alone achieve state-of-the-art performance on variant classification and do so while avoiding the issues that typically impact supervised methods. The benefits of this include better generalization guarantees and providing a source of evidence which is independent and hence complementary to other large-scale efforts, such as population data from biobanks. When compared to some high-throughput functional assays, we found our approach typically achieved equal, if not better performance. This suggests that our method could be used for classification in the same way that some experiments like MAVEs are used when they are relevant assays, *i*.*e*. as strong evidence, as in the case of those performed on BRCA1, P53 and MSH2. While it has long been appreciated that looking at the patterns of protein sequence conservation across organisms can yield insights into disease causing variants in humans, it seems that by bringing together recent developments in machine learning with the rapidly increasing amount of sequencing data from diverse organisms, we can extract more precise statements than previously realized and on a sufficiently large scale to be able to impact our sum knowledge of the clinical significance of disease causing variants. The present model is dependent on the quality of multiple sequences alignment, however recent progress towards alignment-free protein language modeling may help bridge that gap in the future^53,54^. Our data and results, available at evemodel.org, provide information on a gene-by-gene basis where researchers and physicians can look at individual variants in detail, including scores and charts enabling assessment of appropriate tradeoffs between sensitivity and specificity in the context of a particular disease (Fig. 5). We also provide training data and benchmark sets for researchers wishing to develop their own models.

**Figure 5.**
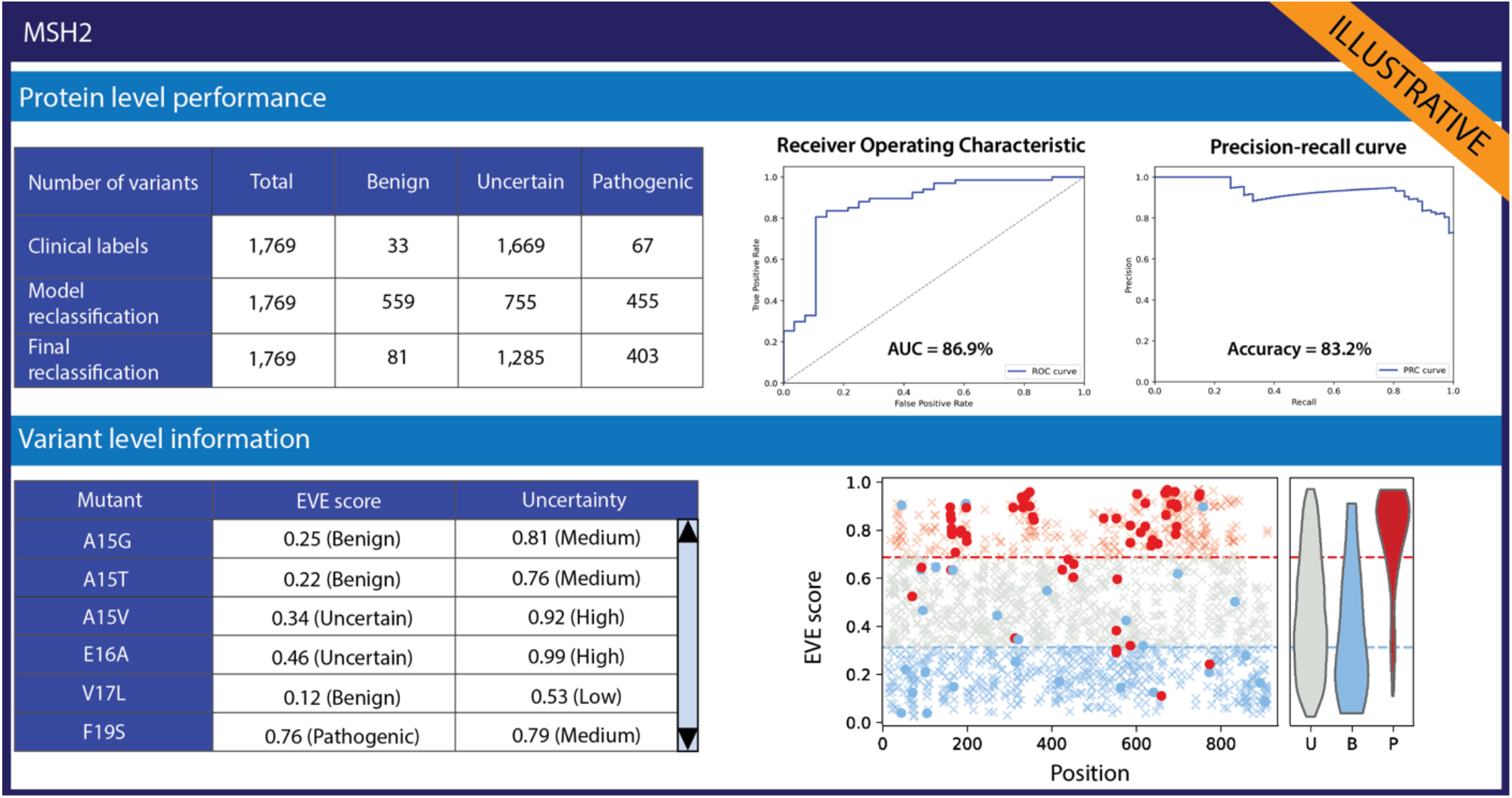
EVE protein profile card (MSH2). Profile cards provide key information to practitioners about each protein: aggregate AUC/Accuracy, performance curves (ROC & Precision-recall) and variant-level EVE scores & uncertainties. Source data for all protein curves & predictions is available in supplementary material.

Despite the usefulness of labelling variants as pathogenic or benign, the reality of clinical assessment is often much more nuanced. Different variants of the same gene, and even the same variant, can lead to vastly different disease severity or even different diseases, aspects masked by the use of simple discrete pathology categories. Whilst the continuous EVE score, as opposed to the discrete classifications, may be used in specific cases to assess disease severity (as with SCN5A), there is not enough clinical data to validate this globally.

Although we do not explore combinations of variants in this work, our model should be sufficiently powerful to address dependencies as the Bayesian VAE computes the evolutionary index of a variant in the context of all other residues in the protein. Our initial exploration suggests that humans have on average ~12% of their genes with two or more variants compared to a reference genome and that for the 59 “ACMG actionable genes” there are ~21k combinations of variants occurring in total ~1.5 million times (estimated using 50,000 whole exomes in the UKBioBank^1^) (Methods, Extended Data Fig. 8, Supplementary Table 8).

The primary advantage of our approach over experimental approaches is significant gain in scope at a negligible fraction of the cost. An appealing prospect is that our methods may be useful in guiding future experimental efforts, essentially acting as a means of identifying which variants and in which genes it would be most informative to probe and the prioritization of genes where evolutionary information is scarce or the quality of the sequence alignment is poor.

We conclude with a remark on biodiversity. Our models are trained on sequences from diverse organisms. The 1,081 models we have trained to date make use of data from over 139k organisms. Of these, we identified 17k organisms which are on the International Union for Conservation of Nature’s (IUCN) Red List of Threatened Species^55^ including 1,301 classified as vulnerable, 1,148 endangered, 548 critically endangered, 10 extinct in the wild and 21 extinct organisms. Our analysis is one small but unusually direct demonstration of how the biodiversity of life on Earth benefits human health. Models designed to aid clinical decisions are improved by incorporating information from more diverse organisms.

## Methods

### Data acquisition

#### Choosing subset of clinically relevant proteins and building multiple sequence alignments from UniRef100 for model training

In this study, we sought a list of ~1, 000 genes associated with disease. We selected our ~1, 000 subset by searching ClinVar for genes with at least 1 Benign and 1 Pathogenic label (detailed below), and then prioritised genes based on the number of high-quality labels available, as well as compute resources required to obtain multiple sequence alignments and infer parameters of the the Bayesian VAE for that protein (*e*.*g*. we avoided extremely large genes such as TTN. To each gene we associate a single protein, by picking the canonical transcript according to Uniprot/Swissprot [1]. To train EVE, we build multiple sequence alignments for each protein family by performing five search iterations of the profile HMM homology search tool hmmer [2] against the UniRef100 database of nonredundant protein sequences [3]. Following the protocol of Hopf et al. [4] and Riesselman et al. [5], for each alignment we only keep sequences that align to at least 50% of the target protein sequence, and columns with at least 70% residue occupancy.

We explore a range of bit scores, using 0.3 bits per residue as a reference, and select the best possible multiple sequence alignment based on the criteria of maximal coverage of the target protein sequence and sufficient, but not excessive, number of sequences in the alignment (the latter implying an alignment that is too lenient). Specifically, we prioritize alignments with coverage *L*_cov_ ≥ 0.8*L*, where *L* is the length of the target protein sequence, and with a total number of sequences *N* such that 100, 000 ≥ *N≥* 10*L*. If these requirements cannot be met, we sequentially relax them down to *L*_cov_ ≥ 0.7*L* and *N≤* 200, 000 until we find the best available multiple sequence alignment for each protein. Following this procedure, we obtain a set of 1,081 clinically relevant proteins with corresponding evolutionary training data (Supplementary Table 1).

#### Obtaining clinical significance labels from ClinVar

To obtain variants with clinical labels we make use of the ClinVar public archive [6], which contains reports of the relationships between human genetic variation and phenotypes, with supporting evidence. Of particular relevance for this work is the ACMG–AMP consistent classification of single nucleotide variants (SNVs) into five categories: Benign, Likely Benign, Uncertain Significance, Likely Pathogenic and Pathogenic. In addition, the quality of evidence provided is ranked according to a four star system, which can be summarized as follows:

- No Stars – Interpretation provided but criteria not met.
- One Star – Criteria provided, single submitter.
- Two stars – Criteria provided, multiple submitters, no conflicts.
- Three Stars – Reviewed by expert panel.
- Four Stars – Practice guideline.

Of ~ 78k missense variants labeled (Likely) Benign or (Likely) Pathogenic in ClinVar, ~ 63k have one star or more. In most of this work, with the exception of Extended Data Fig. 6, we only consider labels with quality rating of one star or higher.

While ClinVar contains clinical labels of SNVs, our model provides evidence at the amino acid variant level. We therefore require a procedure that selects a single label whenever more than one SNV is present in ClinVar for the same amino acid substitution. For these cases we pick the label with the most stars, and if two SNVs have labels with equal star-rating, we pick the most recent.

Finally, in order to obtain a high quality set of labels for validation, we assign all no-star labels as Uncertain and collapse the remaining (Likely) Benign and (Likely) Pathogenic into just two classes, Benign and Pathogenic, respectively. Unless stated otherwise, this is the set of benign and pathogenic labels used for benchmarking throughout the text. In total, we have 25k such labels across 1,081 proteins. Beside all variants labeled with Uncertain Significance and with no-star rating in ClinVar, we also define as Uncertain all variants observed in gnomAD (detailed below) which do not feature in ClinVar. In total we find 363k “variants of unknown significance” across the 1,081 explored in this work.

For the purpose of comparison to high-throughput experiments, we find the above label definitions too restrictive, with only 5 genes having both a substantial number of these high quality labels, as well as high-throughput experimental data (Supplementary Table 5). To expand our analysis to include more experiments, we therefor define a second more lenient label policy. We define “Lenient” labels as the set of all labels in Clinvar – including no-star rating ones – as well as defining as Benign all variants which are more frequent in the population than the most frequent Pathogenic label in the same gene (frequencies estimated from gnomAD, see below). This policy is similar to the one used in Refs [7, 8]. It is important to stress that we expect these labels to be less trustworthy. Consistent with this, our model performance improves as we consider sets of labels with more stringent quality controls – our average AUC over all 1,081 proteins improves from 0.86 to 0.92 to 0.93 as we compute it against ClinVar lenient, 1-star or higher and 2-star or higher labels, respectively (Extended Data Fig. 6a).

#### Population data from gnomAD

The Genome Aggregation Database (gnomAD) [9], seeks to aggregate exome and genome sequencing data from a variety of large-scale sequencing projects, and provide summary data. We make use of both the v2 and v3 data sets. The v3 data set spans 71,702 genomes from unrelated individuals sequenced as part of various disease-specific and population genetic studies. The v2 data set spans 125,748 exomes and 15,708 genomes from unrelated individuals, again, sequenced as part of various disease-specific and population genetic studies, totalling 141,456 individuals. The gnomAD coalition removes individuals known to be affected by severe pediatric disease, as well as their first-degree relatives, however some individuals with severe disease are potentially included in the data sets, albeit likely at a frequency equivalent to or lower than that seen in the general population [9].

Our downstream analysis makes use of this data for estimating frequency of amino acid variation over the population. For this purpose, we take amino acid variant frequencies to be the sum of frequencies of all missense SNVs coding for the same amino acid substitution.

#### Population data from UKBiobank

The UKBiobank [10] is an unprecedented large-scale biomedical database containing in-depth genetic and health information from half a million UK individuals. While it contains health-related records, bio-markers and detailed information about the lifestyle of the participants, in this work we only make use of the genomics aspect of this resource. In particular, we use the 50k whole exome sequencing cohort release from February 2020 assembled using a corrected SPB pipeline that converts raw sequencing data to a quality-controlled set of population variation. Unlike gnomAD which only provides population wide summary data, UKBiobank provides all SNVs for each individual genome, so we use this data to estimate the overall prevalence of more than one variant per gene across the population (Extended Data Fig 8). We also use this data to extract all seen pairs of amino acid variants in the same gene, and their frequencies, across the 59 “actionable genes” as defined by ACMG [11] (Supplementary Table 8).

### EVE Modeling approach

#### Limitations of supervised modeling methods

Computational methods for variant effect classification so far have formulated the problem as a supervised learning task, training models to discriminate between pathogenic and benign labelled examples. These methods are however plagued by several biases:

- **Noisy labels:** Protein variant labelling in ClinVar is an error prone process [12]. When the number of available labels is too small to properly infer the scale of noise, supervised models risk overfitting the training data, leading to poor generalization performance [13, 14];
- **Selection bias:** The distribution of labels is not uniform across proteins. Pathogenicity is further studied for some proteins than others, with some proteins having no clinical label at all. Labelling may be scarce for variants coming from under-represented groups, and more common for others groups (e.g., European ancestry). Supervised models trained on such biased datasets may subsequently infer incorrect statistical relationships between covariates and labels, and thus generalize poorly [15];
- **Class imbalance:** The proportion of pathogenic labels in the data does not reflect the natural frequency of pathogenic variants, in part as a consequence of the criteria governing the class assignment (less evidence is required for a variant to be labelled ‘pathogenic’ Vs ‘benign’)[12]. Not rebalancing the training data will lead to biased probability estimates for the different classes. However, rebalancing may lead to noise misestimation, and may be challenging to do properly in practice [16];
- **Data leakage:** Reported performance may be inflated due to non-obvious data leakages and improper cross-validation splits (e.g., prediction accuracy at a given position may be inflated if the model had access to labels at that position during training). [13, 14, 17]

Given these significant limitations, we set out to instead tackle the problem in a *fully unsupervised* manner: labels are only used during final model evaluation, not during model training.

#### A fully unsupervised approach to the variant effect prediction task

The natural distribution of protein sequences we observe across organisms is the result of billions of evolutionary experiments. By modeling that distribution, we implicitly capture constraints on the protein sequences that maintain fitness [4, 5]. We hypothesize that this distribution may be used to estimate the fitness change for any variant or combination thereof, even for protein sequences that have not been observed. By comparing relative likelihoods of different sequences, we may in turn be able to set apart the sequences that are likely pathogenic.

#### Learning distributions over protein spaces

Variational Autoencoders (VAEs) [18, 19] have been shown to be effective at learning complex high-dimensional distributions across a wide range of tasks – from computer vision, to natural language processing [20], to molecules modeling [21], and many others. In this work, we train VAEs to infer a distribution over amino-acid sequences for each protein. More formally, for a given protein family *p*, we learn a distribution *p*(**s**| *θ* _**p**_) where **s** is a fixed-length amino-acid sequence and *θ* _**p**_ are the model parameters for that protein family. Variational Autoencoders make the assumption that the data **s** are generated from a latent variable **z**, and model the conditional distribution *p*(**s**| **z**, *θ* _**p**_) with a neural network architecture (the “decoder”, with parameters *θ* _**p**_). We use amortized inference and model the approximate posterior distribution *q*(**z**| **s**, *ϕ*_**p**_) with a neural network as well (the “encoder”, with parameters *ϕ*_**p**_). Similarly to Riesselman et al. [5], we obtain stronger performance with a Bayesian VAE in which we learn a fully-factorized Gaussian distribution over the decoder weights *θ* _**p**_. We interpret this observation by the fact that in a VAE architecture the encoder is trained only in-distribution and will extrapolate arbitrarily when dealing with out-of-distribution points. Thus, for mutants far from the training data, latent positions obtained from the encoder will be less reliable. While a standard VAE will then decode that mutant to an arbitrary position, the output from a Bayesian VAE will average over decodings (the decoder being a Bayesian Neural Network), which will dampen the corresponding probability estimates over amino acid positions [22].

#### Training VAEs over Multiple Sequence Alignments

The distribution of protein sequences available in genomic databases is biased by human sampling (e.g., certain species of interest are more sequenced than others) and evolutionary sampling (phylogeny). Not correcting for these biases in the Multiple Sequence Alignments (MSAs) that we extract from genomic databases to train our models will lead to learning an improper probability distribution. Following the approach described in Ekeberg et al. [23], we correct for these two biases by re-weighting each protein sequence **s**_*i*_ from a given MSA according to the reciprocal of the number of sequences in the corresponding MSA within a given Hamming distance cutoff *T*.

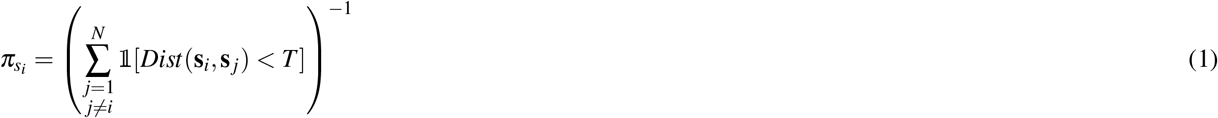

where *N* is the number of sequences in the MSA, *i* indexes over proteins, and **s** _*j*_ are other protein sequences in the MSA. Similarly to Hopf et al. [4], we set *T* = 0.2 for all human proteins. During model training, we sample each mini-batch element by sampling sequences according to their weight *π*_*s*_.

We train the VAE models by maximizing the Evidence Lower Bound (ELBO) which forms a tractable lower bound to the log-marginal likelihood. Following the “Full Variational Bayesian” approach described in [18], the ELBO is expressed as follows:

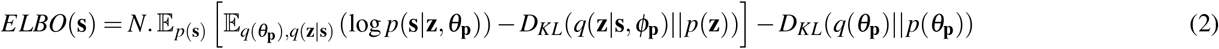

where N is the size of the training data and *p*(*z*) is the prior distribution over latent variable *z* (standard Gaussian), and *D*_*KL*_ is the Kullback–Leibler divergence (a measure of dissimilarity between two probability distributions). In our Bayesian VAE formulation, we learn a fully-factorized Gaussian distribution *q*(*θ* _**p**_) over the decoder parameters, with standard Gaussian prior *p*(*θ* _**p**_). Similar to Riesselman et al. [5] and in agreement with the sequence re-weighting scheme, we set 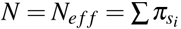, the effective number of sequences in the MSA defined as the sum of the different sequence weights.

#### Model architecture

We performed several ablation analyses to optimize the underlying model architecture and the choice of training hyperparameters (see Extended data Fig. 1). The main changes we suggest over the DeepSequence architecture [5] are as follows:

- Symmetrization of the encoder and decoder network architectures;
- Increased number of layers and increased layer width for the encoder and decoder networks (2, 000 - 1, 000 - 300 and 300 - 1, 000 - 2, 000 respectively);
- Increased size of the latent space (50);
- Larger number of training steps to train the more complex architecture (400, 000 training steps);
- Lower learning rate to stabilize learning process (10^*-*4^);
- Removal of the group sparsity priors, responsible of significant performance drops for certain proteins;

Extended Data Fig. 2 summarizes the performance gains achieved with the changes above compared with DeepSequence, by comparing Spearman correlation of the two models with the output of 38 different benchmark MAVEs, following the same protocol as described in Riesselman et al. [5].

#### Evolutionary index

We define the *evolutionary index* of a protein variant **s** as the relative fitness of **s** compared with that of a wild-type sequence **w**. Building from the probabilistic viewpoint that gave rise to the Bayesian VAE, fitness of a sequence may be measured by the difference in log-likelihood of that sequence with the wild type. Since estimating the exact log-likelihood is intractable in practice, we use the negative ELBO, a bound on the log marginal likelihood, as an approximation which leads to strong empirical results (Fig. 2). The *evolutionary index E*_*s*_ of protein **s** may therefore be expressed in terms of a tractable difference between two ELBOs:

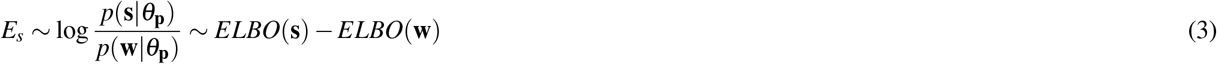

For each variant **s** of interest, we estimate ELBO values by averaging over 200k samples from the approximate posterior distribution *q*(**z**|**s**, *ϕ*_**p**_).

### Separating pathogenic and benign variants with probabilistic clustering

*Evolutionary indices* show very strong correlations with existing clinical labels across proteins (Fig. 2a). We fit a Gaussian Mixture model with two components directly on the evolutionary index distribution, in order to automatically separate variants into Pathogenic and Benign clusters. Besides the advantage of being fully unsupervised, performing probabilistic clustering of variants allows us to quantify our uncertainty about the class assignment (see next section).

The Gaussian Mixture Model (GMM) [24] is a probabilistic model that assumes the data are generated from a mixture of a finite number of Gaussian distributions with unknown parameters. We experiment with different architectures and training algorithm: single model Vs hierarchical, training with Variational Inference (VI) Vs Expectation-Maximization (EM) algorithm. We obtain the best performance on the downstream task with the following approach. We first train an overarching two-component GMM on the distribution of evolutionary indices for all single amino acid variants for all 1,081 proteins combined. We use the resulting parameters to initialize the Gaussians of the protein-specific GMMs, fit on all single amino acid variants for each protein separately. Intuitively, the cluster with higher mean will contain sequences with higher evolutionary indices, and therefore less likely sequences under the learnt sequence distribution. For each model after convergence, we thus define the component with the highest mean as the Pathogenic cluster, and the other one as the Benign cluster. We then form a global-local mixture of GMMs to combine the cluster predictions from the overarching GMM with the predictions from the protein-specific GMM for each protein. More formally, for a given protein **s** with evolutionary index **E**_**s**_, from a protein family p, we have:

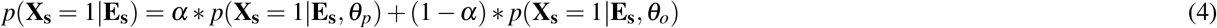

where *X*_*s*_ is a binary random variable equal to 1 if **s** is pathogenic (0 if benign), *α* represents the relative weight of protein-level GMM in the ensemble (set to 0.3 via grid search with respect to average accuracy and AUC), *θ* _*o*_ and *θ* _*p*_ are the parameters of the overarching GMM and protein-specific GMM respectively. *p*(**X**_**s**_ = 1|*z* **E**_**s**_) is what we refer to as the *EVE score* which quantifies the propensity of a given variant to be pathogenic.

#### Quantifying the uncertainty in the cluster assignment

A crucial benefit of a probabilistic clustering approach is the ability to identify the set of variants for which the classification is the most uncertain. We measure the total uncertainty on the cluster assignment for a protein **s** via the corresponding predictive entropy PE:

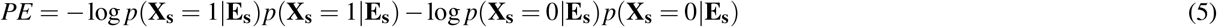

As we increase the proportion of variants excluded based on their predictive entropy (excluding higher values first), the AUC/accuracy monotonically increases (Fig. 2b and 2c). This confirms the ability of the predictive entropy metric to properly identify uncertain variants. This property allows us to set aside an increasing proportion of variants for which we are the most uncertain about, thereby reaching a desired level of average accuracy on the variants that we do classify. In our large scale classification following ACMG-AMP guidelines, we exclude the 25% of variants across the 1,081 proteins that had the highest uncertainty, leaving us with variants with an expected accuracy greater than 90% based on available clinical labels.

### Evaluation

#### Overall validation of EVE with clinical labels

We measure the aggregate performance of EVE models in terms of their ability to properly discriminate between pathogenic and benign variants for which clinical labels already exist. As discussed above, we focus on the subset of highest quality labels for the purpose of model evaluation. Aggregate performance is measured via the area under the Receiver Operating Characteristic curve (AUC) and the total prediction accuracy (setting a threshold that equally balances out false positives and false negatives). Our main results are summarized in Fig. 2b, 2c, 2d (see also Supplementary Table 2 for detailed results across all 1,081 proteins). We looked at performance across different proteins groups: the full set of 1,081 proteins and the subset thereof with 3, 5 and 10 Benign and 3, 5 and 10 Pathogenic labels - denoted as 3B3P, 5B5P and 10B10P, respectively. The overall AUC and accuracy are high across the board and across subsets of proteins considered (all AUCs are above 0.9, and all accuracies above 82%). Performance is even higher on the subset of variants for which the model is the most confident about (e.g., accuracy above 98% and AUC above 0.98 on the set of proteins in the bottom uncertainty decile).

Additionally, we provide the ROC and Precision-recall curves for all proteins, on the full set of variants and on the set where we exclude the 25% most uncertain variants (available on the different protein profile cards, e.g. Fig. 5, Extended data Fig. 4 and supplementary material). These may help practitioners to make more informed decisions based on the relative importance of false positives and false negatives characterizing their use case.

#### Benchmarking EVE with supervised models of variant effects

We compare the performance of EVE models with that of the top performing supervised variant effect predictors (VEPs) published to date: M-CAP [25], MISTIC [26], and Polyphen [27] (Extended data Fig. 4). Since we do not account for the train-validation split used by these models at train time (as some of these methods did not make this data publicly available), the reported performance for VEPs is to be interpreted as an upper bound on their true performance (due to potential data leakage). When comparing methods, we compute AUCs or accuracy using the high quality labels defined above, for the intersection of variants for which the models have made predictions. EVE models outperforms all VEPs when considering the largest set of variants possible.

#### Benchmarking against high-throughput functional assays

We compared the performance of EVE to a number high-throughput experiments [7, 28, 29, 30, 31, 32, 33, 34, 35, 36], which were also designed with the intention of predicting the pathogenicity of variants. Our approach to comparing methods, follows much the same process as when comparing EVE to VEPs using AUCs – again we consider only the intersecting set of variants common to both the assay and EVE when computing the AUC. One difference however is that we perform the comparison using multiple label definitions as described above, as well as older versions of ClinVar, to avoid comparing EVE and the experiment on labels which were established by making use of data from that same experiment. On average we find EVE outperforms the experiments regardless of this choice (Supplementary Table 5).

In Figure 3 we use the high quality labels, distinguishing ClinVar 2020 data (lighter red, lighter blue) from ClinVar 2017 (darker red, darker blue), when the experiment has been used in establishing labels in the 2020 data. In particular, for both model and experiment, we report AUCs for P53 and BRCA1 using 2017 ClinVar data, and for PTEN, SCN5A and MSH2 usinf 2020 ClinVar data.

In Extended Data Figure 6b we use “lenient” labels (defined in Data Acquisition section), selecting 2020 or 2017 ClinVar data as appropriate and noted in (Supplementary Table 5).

### Combining EVE predictions with other sources of evidence for variant classification

The 2015 American College of Medical Genetics and Genomics–Association for Molecular Pathology (ACMG-AMP) guidelines [12] present steps towards a systematic approach to variant classification which can be used consistently across independent groups. They propose a classification scheme consisting of 28 criteria to classify variants into one of five categories – Benign, Likely Benign, Uncertain, Likely Pathogenic and Pathogenic. Each criteria corresponds to a different form of evidence, such as population data, or functional data, and the strength of evidence provided by a given criteria falls into one of four categories – Supporting, Moderate, Strong and Very Strong. Finally, a set of rules determines how criteria are to be combined to determine the category of a given variant. For instance, one Strong pathogenic criterion, combined with two or more Supporting criteria, would result in a variant being classified as Likely Pathogenic.

One of the defining characteristics of our model is the fact that it only uses one source of evidence – evolutionary data – to score variants according to their clinical significance. As such, it is straightforward to combine the EVE scores with other evidence in a manner consistent with the strategy outlined by the ACMG-AMP. We have argued that our approach yields evidence that can be of equal or superior quality to a functional assay, and as such, should be regarded as Strong evidence in these cases. We now detail how we collect different lines of evidence and implement them in our analysis to classify as many variants as possible. We refer to this process as *large N variant classification* to distinguish considerations relating to the need to classify at scale from considerations relating to the classification of individual variants.

In practice, only a small number of the other ACMG-AMP criteria are amenable to use in large *N* variant classification at present. Of the 28 criteria defined by ACMG-AMP, we only make use of 4 in our analysis:

- **Strong Benign criterion 1 (***BS*1**) Definition**: Allele frequency is greater than expected for disorder.
- *BS*1 **Implementation**: We make use of population data from gnomAD [9]. For a variant to satisfy this criterion, we require that it be observed more frequently in gnomAD than any known (Likely) Pathogenic variant in that gene. In other words, we require a lower bound on the frequency *ν*_var_ *>* max(*ν*_path_).
- **Supporting Benign criterion 6 (***BP*6**) Definition**: Reputable source recently reports variant as benign, but the evidence is not available to the laboratory to perform an independent evaluation.
- *BP*6 **Implementation**: We took variants in ClinVar labelled as Benign but with a zero-star rating to satisfy this criterion.
- **Moderate Pathogenic criterion 2 (***PM*2**) Definition**: Absent from controls (or at extremely low frequency if recessive) in Exome Sequencing Project, 1000 Genomes Project, or Exome Aggregation Consortium.
- *PM*2 **Implementation**: This criterion is the pathogenic equivalent of *BS*1. Again we use frequency data from gnomAD and this time we require that for a variant to satisfy this criteria its frequency must be lower than any known (Likely) Benign variant in that gene, *ν*_*var*_ *<* min(*ν*_benign_).
- **Moderate Pathogenic criterion 5 (***PM*5**) Definition**: Novel missense change at an amino acid residue where a different missense change determined to be pathogenic has been seen before.
- *PM*5 **Implementation**: Any variant found at the same position as a (Likely) Pathogenic variant with at least a one-star rating in ClinVar met this criterion.

### Combining criteria

If we take our models as capable of providing strong evidence of a variant being either benign *BS*_EVE_, or pathogenic *PS*_EVE_ then there are 4 ways in which our model evidence can be combined with the above criteria to reclassify a VUS as Benign, Likely Benign, or Likely Pathogenic:

- **Benign**: *BS*_EVE_ and *BS*1
- **Likely Benign**: *BS*_EVE_ and *BP*6
- **Likely Pathogenic**: (*PS*_EVE_ and *PM*2), or (*PS*_EVE_ and *PM*5)

Whenever the conclusions of this analysis are conflicting with an existing ClinVar assignment, we set the variant as Uncertain. We provide both summary statistics (Fig. 4d. Supplementary Table 6) as well as a complete list of evidence used for the classification of every variant in our analysis (see data availability).

#### Comments on criteria not used in our analysis

There are additional criteria in principle suitable for large *N* variant classification that we did not make use of due to concerns of “double counting” particular forms of evidence. Broadly, this problem is well known [37] but also manifests in somewhat unique ways in our analysis. As such, here we comment on how these issues would arise in our analysis should we try to make use of certain criteria.

- **Moderate Pathogenic criterion 1 (***PM*1**) Definition**: Located in a mutational hot spot and/or critical and well-established functional domain (*e*.*g*., active site of an enzyme) without benign variation.
- *PM*1 **Discussion**: EVE, being a model for all possible amino acid variants in a protein sequence, is a natural hot/cold spot detector. The objective of the classification exercise, however, is to combine distinct sources of evidence, so we leave this criterion out of our analysis.
- **Supporting Benign criterion 4 (***BP*4**) Definition**: Multiple lines of computational evidence suggest no impact on gene or gene product (conservation, evolutionary, splicing impact, etc.)
- **Supporting Pathogenic criterion 3 (***PP*3**) Definition**: Multiple lines of computational evidence support a deleterious effect on the gene or gene product (conservation, evolutionary, splicing impact, etc.)
- *BP*4 **and** *PP*3 **Discussion**: The model we propose in this paper should in principle fall into this category but we have argued that by using an unsupervised approach it does not suffer from label biases and it can be subjected to more stringent validation. This results in a more reliable form of evidence than is provided by current state-of-the-art computational approaches, and we argue that it can stand alone as a source of strong evidence. We could in principle use other computational methods as an additional source of evidence in our classification pipeline, however this does not seem reasonable since almost all computational methods make at least some use of evolutionary data and population frequency data, both of which have already been made use of. We therefore opt for leaving this criterion out of our analysis.
- **Supporting Pathogenic criterion 5 (***PP*5**) Definition**: Reputable source recently reports variant as pathogenic, but the evidence is not available to the laboratory to perform an independent evaluation.
- *PP*5 **Discussion**: In direct analogy to BP6, we could take variants with zero-star rating Pathogenic labels in ClinVar to satisfy this criterion. In practice this would not alter our results, since according to ACMG–AMP guidelines a single Supporting Pathogenic criterion is not sufficient to impact variant classification.

### Identifying potentially incorrect labels

A benefit of our approach being unsupervised is that it provides a natural means of identifying potentially incorrect labels. While there was very good agreement between our models and the vast majority of clinical labels, there were a small number of variants for which there was strong disagreement. There are three possible reasons for these disagreements

1. The model failed in some way (*e*.*g*. the model was trained on a poor quality multiple sequence alignment or the training process itself had a problem.)
2. Evolutionary data does not reflect the clinical significance in these special cases.
3. The clinical label is incorrect.

In some cases, we can flag variants for which the third is the potential cause of disagreement by looking for consensus with other sources of evidence. As other sources of evidence, we used the ones described in the previous section. We identified 164 such variants.

### Data availability

The data analysed and generated in this study is available at www.evemodel.org.

## Notes

http://evemodel.org/

### Competing Interest Statement

The authors have declared no competing interest.

