## Extended Data Figures 1-8 and Table 1 for "Large-scale clinical interpretation of genetic variants using evolutionary data and deep learning"

Extended Data Figure 1

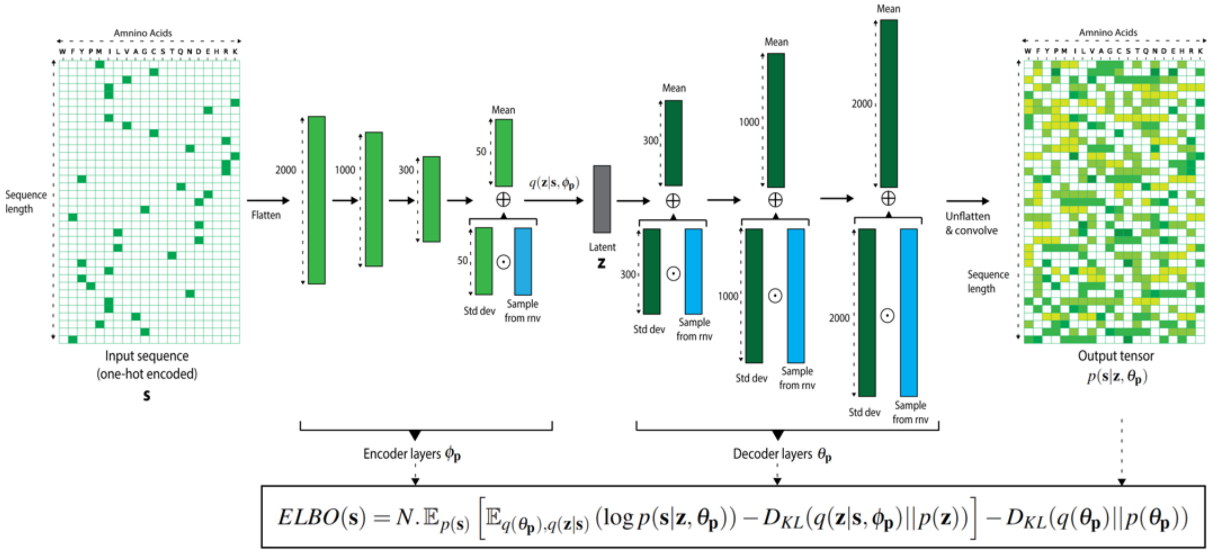


**Extended Data Figure 1 – Bayesian VAE architecture details.** The Bayesian VAE architecture in EVE is comprised of a symmetric 3-layer encoder & decoder architecture (with 2,000-1,000-300 and 300-1,000-2,000 units respectively) and a latent space of dimension 50. After performing a one-hot encoding of the input sequence across amino acids (zeros in white, ones in green), we flatten the input before performing the forward pass through the network. We use a single set of parameters for the encoder ($\phi_\mathrm{p}$) and learn a fully-factorized gaussian distribution over the weights of the decoder ($\theta_\mathrm{p}$): weight samples for the decoder are obtained by sampling a random normal variable (rnv), multiplying that sample by the standard deviation parameters, and subsequently adding the mean parameters. A one-dimensional convolution is applied on the un-flattened output of the decoder to capture potential correlations between amino-acid usage. Finally, a softmax activation turns the final output into probabilities over amino acids at each position of the sequence (low values in white, high values in dark green). The overall network is trained by maximizing the Evidence Lower Bound (ELBO), which forms a tractable lower bound to the log-marginal likelihood (Methods).

Extended Data Figure 2

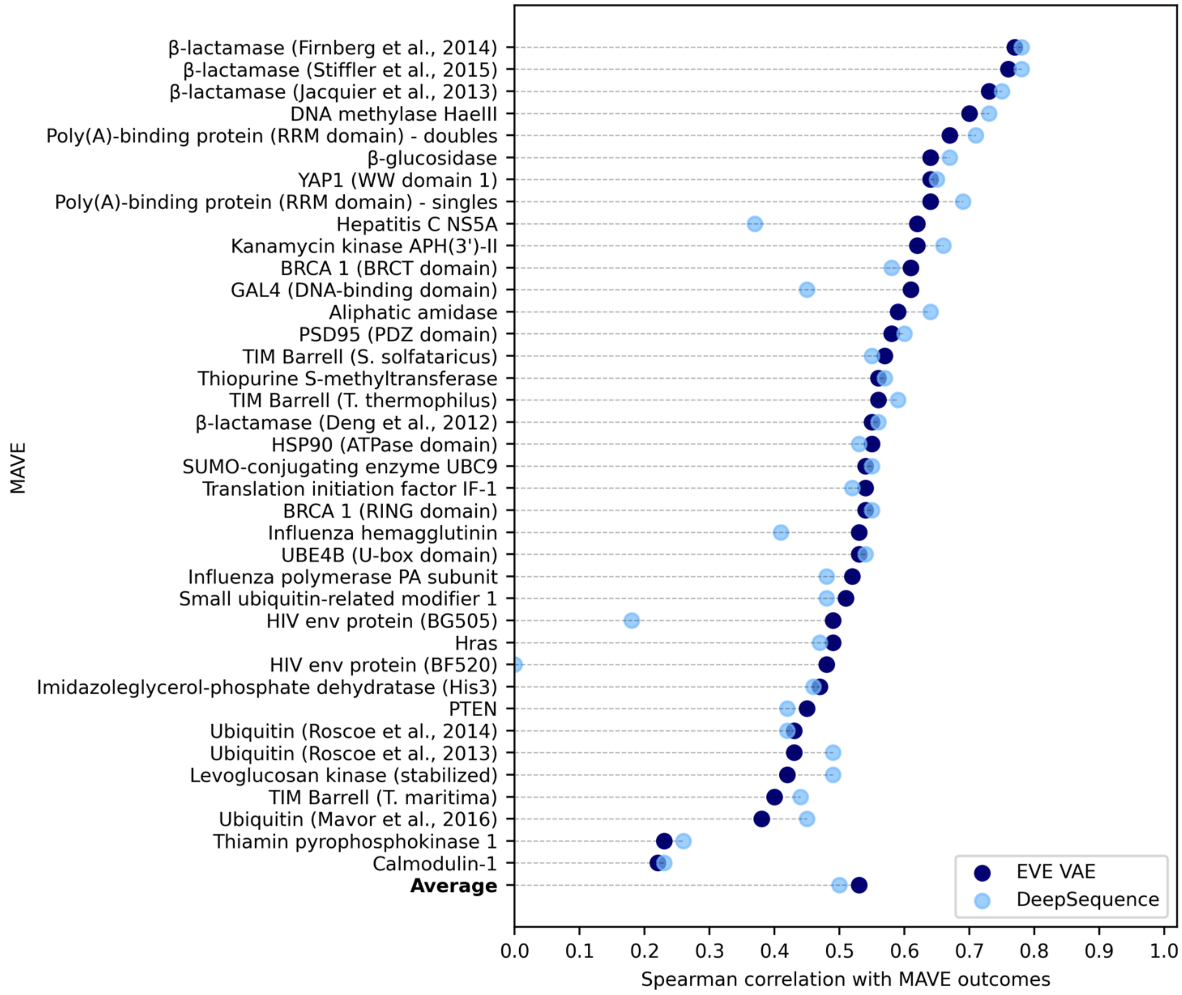


**Extended Data Figure 2 – Comparison of performance at mutation effect prediction of our Bayesian VAE implementation and DeepSequence.** Comparison between the performance of our implementation of Bayesian VAE and the one from state-of-art DeepSequence, at correlating with fitness scores from multiplexed assays of variant effects. The spearman correlations between model and experiment were evaluated for 38 multiplexed assays of variant effects. "Evolutionary indices" were computed using, for both models, the same protocol as Riesselman et al., *i.e.* by sampling 2k times from the approximate posterior distribution (as opposed to the 200k used for reporting EVE results) and by ensembling scores from 5 VAEs.

Extended Data Figure 3

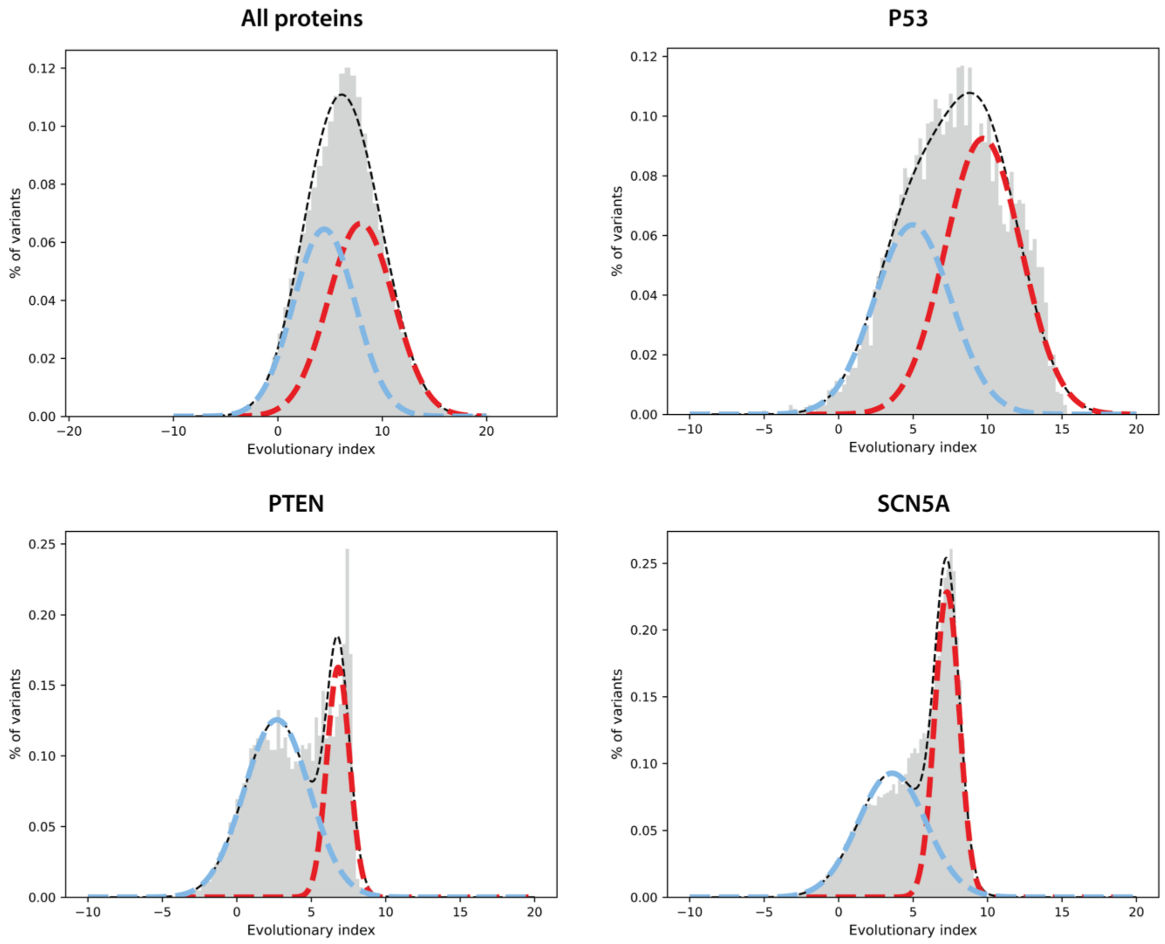


**Extended Data Figure 3 – Evolutionary index distribution overall and by protein.** Two-component Gaussian Mixture Models (GMM) over the distributions of the evolutionary indices for all the single amino acid variants of 1,081 proteins combined (top, left) and for P53, PTEN and SCN5A separately (top right, bottom left and right, respectively). Dashed blue and red lines represent the two components distribution for the benign and pathogenic clusters, respectively, whereas dashed black line represents the GMM distribution.

Extended Data Figure 4

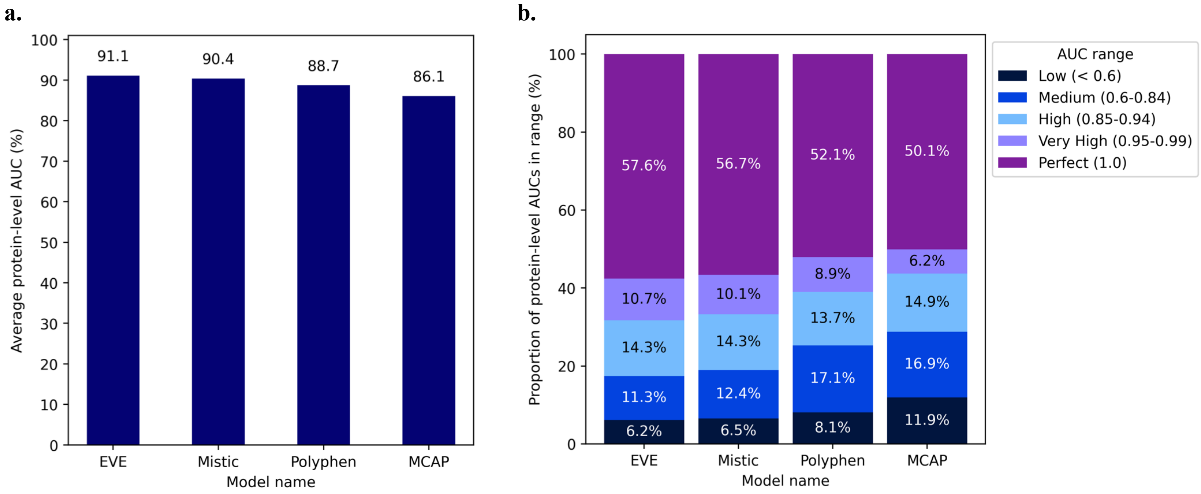


**Extended Data Figure 4 – Comparison of mean AUC of supervised variant effect predictors to EVE.**
**a.** Average AUC per protein, computed across 1,081 proteins, for EVE and state-of-art supervised methods Mistic, MCAP and Polyphen. EVE outperforms existing methods in terms of this metric. **b.** Percentage of the 1,081 proteins for which the model obtains an AUC of less than 0.6, 0.6-0.84, 0.85-0.94, 0.95-0.99 and 1, respectively, for the four models EVE, Mistic, MCAP and Polyhen. EVE has a higher proportion of very high/perfect protein-level AUCs (≥0.95) and a lower proportion of low & medium protein-level AUCs (<0.85) (Methods).

Extended Data Figure 5

**a.**

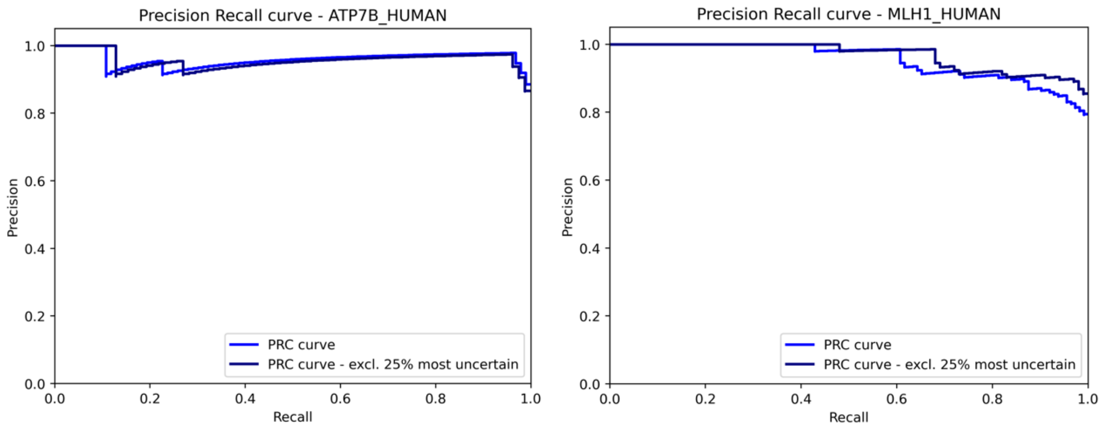


**b.**

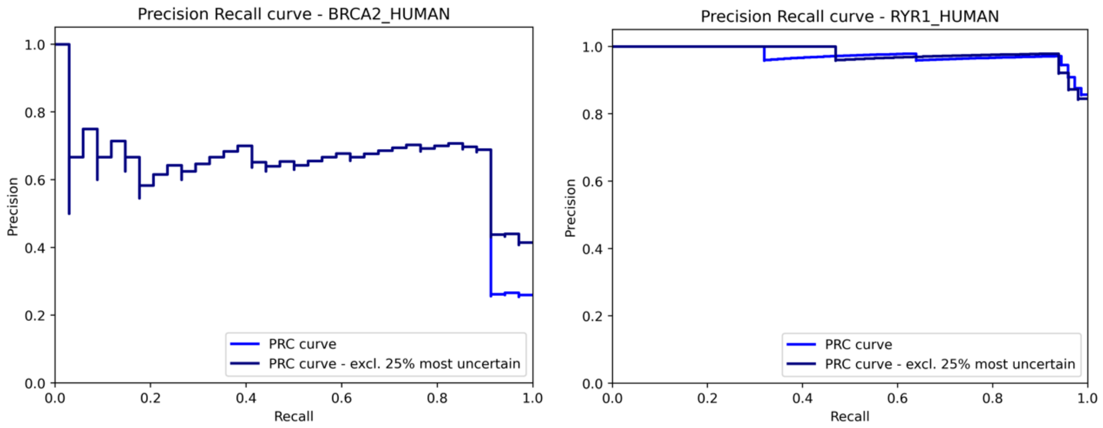


**Extended Data Figure 5 – Comparison of Precision-Recall curves for proteins with similar AUC.**
**a.** Comparison between the precision-recall curve of EVE for ATP7B and MLH1, which both have an AUC over all variants of ~0.88. While for both proteins precision and recall are near perfect, the precision for MLH1 drops for high sensitivity (recall), while for ATP7B it remains high for near perfect sensitivity. **b**. Comparison between the precision-recall curve of EVE for BRCA2 and RYR1, which both have an AUC over all variants of ~0.91. For BRCA2, the precision and recall are considerably worse than for RYR1.

Extended Data Figure 6

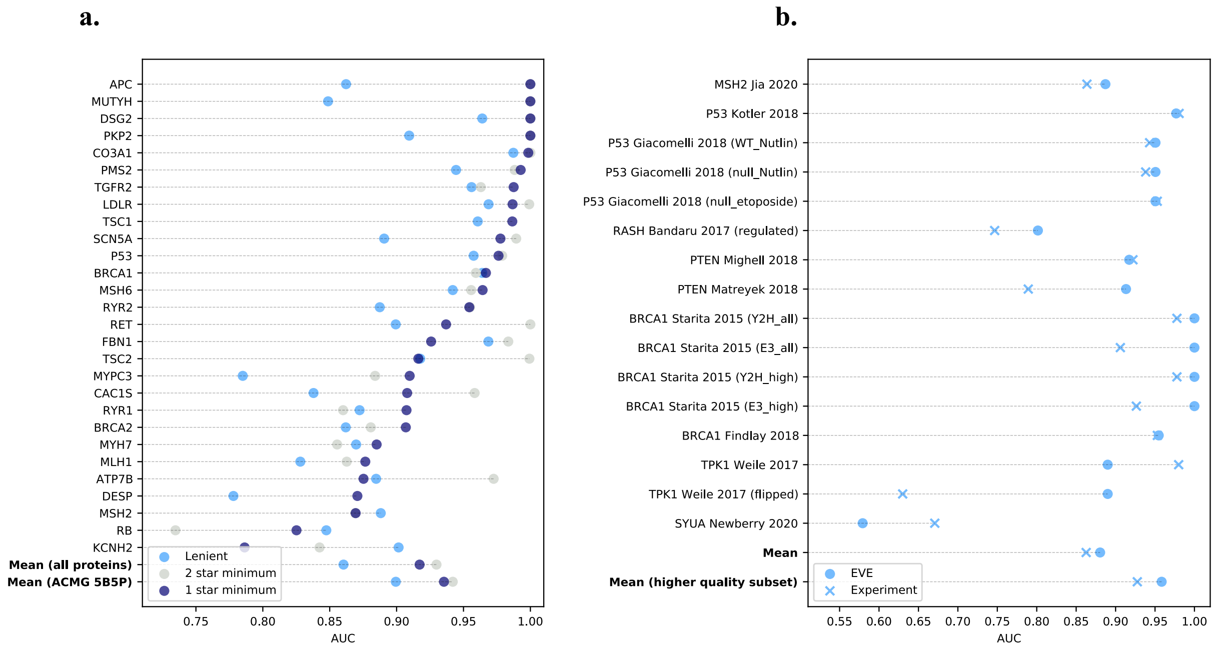


**Extended Data Figure 6. Comparison of label policies, comparison of EVE and experiments. a.** The y-axis is the subset of the ACMG59 actionable protein list with at least 5 benign and 5 pathogenic labels with at least a one-star review status in ClinVar, mean for the 1,081 proteins and mean for this subset. x-axis is AUCs computed using these labels (deep blue), labels with at least a two-star review status (light grey) and a more lenient labelling policy (sky blue), which uses labels from ClinVar of any review status, with additional benign labels added from gnomAD, defined as any variant with a frequency greater than the highest frequency pathogenic variant in that gene (with at least a one-star review status). **b.** AUCs computed with the lenient labels over the set of variant common to both the experimental assay (crosses) and EVE model (dots).

Extended Data Figure 7

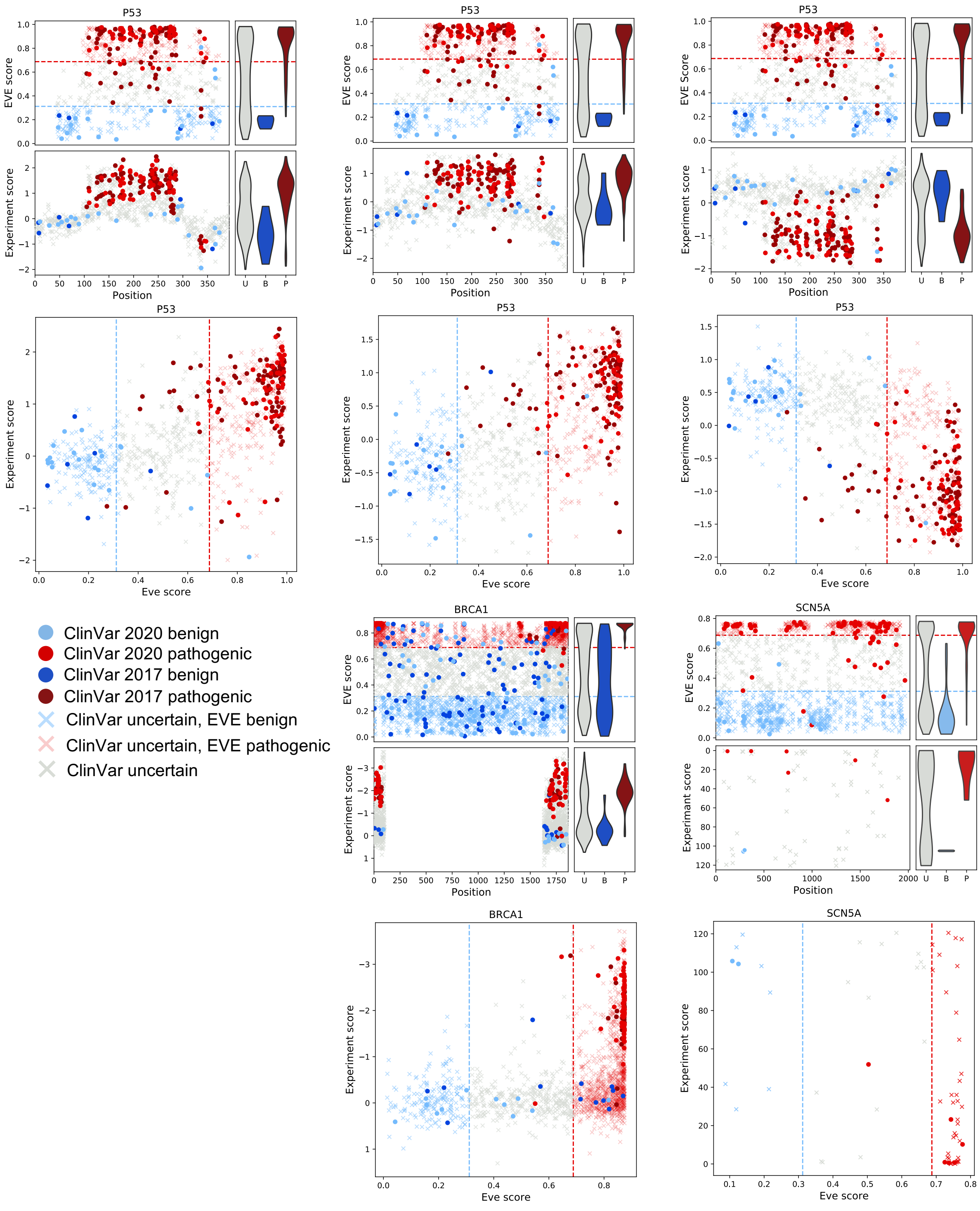


**Extended Data Figure 7. Computational model EVE as good as high-throughput experiments for clinical labels (additional plots).** Comparison of computational model predictions (upper panels, y-axis is EVE score) and experimental assay measurements (lower panels, y-axis experimental assay metric) to ClinVar labels (dots) and VUS (crosses), where pale red and pale blue crosses indicate EVE assignments of VUS. x-axes are position in protein. Also shown are scatter plots of experiment scores (y-axis) against EVE scores (x-axis). Experimental measurements data from deep mutational scans of P53, from left (WT_Nutlin-3, A549_p53NULL_Nutlin-3, A549_p53NULL_Etoposide) SCN5A, and BRCA1.

Extended Data Figure 8

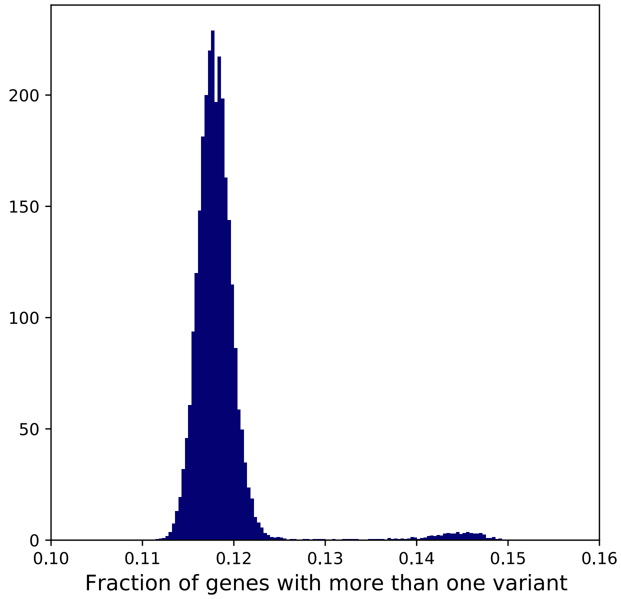


**Extended Data Figure 8 – Fraction of genes per person with more than one variant.** Density function of the fraction of total genes per person (~3k) with at least two variants. Data extracted from 50k genomes of the UKBiobank (Methods).

### Extended Data Table 1

| Protein | AUC | Accuracy 75pct | Labels pre | Labels post | EVE, CV disagree | EVE _and_ACMG, CV disagree | B_LB pre | P_LP pre | U pre | B mod | U mod | P mod | B_LB post | P_LP post | U post |
|---|---|---|---|---|---|---|---|---|---|---|---|---|---|---|---|
| LDLR | 0.99 | 0.98 | 608 | 835 | 17 | 1 | 53 | 555 | 751 | 277 | 319 | 763 | 65 | 770 | 524 |
| FBN1 | 0.93 | 0.97 | 600 | 1166 | 20 | 2 | 16 | 584 | 1981 | 733 | 817 | 1031 | 77 | 1089 | 1415 |
| BRCA1 | 0.97 | 0.83 | 286 | 1795 | 42 | 15 | 176 | 110 | 3925 | 1129 | 1240 | 1842 | 154 | 1641 | 2416 |
| MYH7 | 0.89 | 0.87 | 245 | 586 | 19 | 0 | 11 | 234 | 1355 | 328 | 681 | 591 | 16 | 570 | 1014 |
| TSC2 | 0.92 | 0.84 | 237 | 591 | 23 | 3 | 179 | 58 | 2145 | 678 | 1159 | 545 | 204 | 387 | 1791 |
| BRCA2 | 0.91 | 0.84 | 266 | 1091 | 27 | 5 | 223 | 43 | 5284 | 1533 | 3021 | 996 | 365 | 726 | 4459 |
| MLH1 | 0.88 | 0.88 | 159 | 514 | 15 | 3 | 40 | 119 | 1057 | 351 | 324 | 541 | 53 | 461 | 702 |
| P53 | 0.98 | 0.98 | 153 | 440 | 4 | 1 | 25 | 128 | 737 | 193 | 318 | 379 | 43 | 397 | 450 |
| CO3A1 | 1 | 0.99 | 141 | 477 | 1 | 0 | 19 | 122 | 971 | 416 | 266 | 430 | 83 | 394 | 635 |
| RYR2 | 0.95 | 0.97 | 138 | 766 | 3 | 1 | 28 | 110 | 2695 | 1088 | 1092 | 653 | 275 | 491 | 2067 |
| KCNH2 | 0.79 | 0.91 | 153 | 386 | 11 | 1 | 12 | 141 | 933 | 380 | 397 | 309 | 35 | 351 | 700 |
| ATP7B | 0.88 | 0.9 | 116 | 190 | 9 | 2 | 20 | 96 | 956 | 344 | 439 | 289 | 24 | 166 | 882 |
| SCN5A | 0.98 | 0.96 | 109 | 541 | 4 | 0 | 24 | 85 | 1520 | 665 | 480 | 484 | 114 | 427 | 1088 |
| MSH2 | 0.87 | 0.9 | 100 | 401 | 7 | 2 | 33 | 67 | 1669 | 526 | 886 | 357 | 79 | 322 | 1368 |
| RYR1 | 0.91 | 0.93 | 115 | 605 | 9 | 1 | 25 | 90 | 3359 | 980 | 1939 | 555 | 88 | 517 | 2869 |
| TSC1 | 0.99 | 0.89 | 82 | 198 | 7 | 2 | 77 | 5 | 876 | 453 | 287 | 218 | 70 | 128 | 760 |
| MYPC3 | 0.91 | 0.92 | 73 | 233 | 5 | 0 | 28 | 45 | 1065 | 338 | 528 | 272 | 60 | 173 | 905 |
| MSH6 | 0.96 | 0.95 | 71 | 338 | 2 | 0 | 41 | 30 | 2538 | 1072 | 1178 | 359 | 54 | 284 | 2271 |
| RET | 0.94 | 0.98 | 54 | 161 | 7 | 1 | 13 | 41 | 945 | 536 | 313 | 150 | 25 | 136 | 838 |
| TGFR2 | 0.99 | 1 | 47 | 60 | 0 | 0 | 7 | 40 | 233 | 104 | 75 | 101 | 8 | 52 | 220 |
| CAC1S | 0.91 | 0.76 | 55 | 464 | 9 | 2 | 49 | 6 | 1210 | 508 | 421 | 336 | 240 | 224 | 801 |
| DESP | 0.87 | | 39 | 220 | 0 | 0 | 30 | 9 | 1963 | 648 | 1354 | 0 | 211 | 9 | 1782 |
| RB | 0.83 | 0.84 | 43 | 114 | 5 | 0 | 29 | 14 | 528 | 211 | 292 | 68 | 58 | 56 | 457 |
| DSG2 | 1 | 0.96 | 43 | 158 | 1 | 0 | 37 | 6 | 763 | 240 | 375 | 191 | 60 | 98 | 648 |
| APC | 1 | 1 | 37 | 1655 | 0 | 0 | 29 | 8 | 3828 | 1465 | 1271 | 1129 | 605 | 1050 | 2210 |
| PMS2 | 0.99 | 1 | 37 | 263 | 0 | 0 | 25 | 12 | 1364 | 422 | 665 | 314 | 29 | 234 | 1138 |
| MUTYH | 1 | 0.89 | 20 | 33 | 2 | 0 | 9 | 11 | 738 | 435 | 323 | 0 | 23 | 10 | 725 |
| PKP2 | 1 | 1 | 27 | 139 | 0 | 0 | 20 | 7 | 677 | 259 | 347 | 98 | 61 | 78 | 565 |

**Extended Data Table 1 – Classification summary for actionable genes defined by ACMG with at least 5 benign and 5 pathogenic ClinVar labels.** AUC, accuracy for the most certain variant predictions (dropping the 25% most uncertain). Labels pre: Number of ClinVar labels with at least a one-star rating, Labels post: Number of labels following our analysis combining EVE predictions with other sources of evidence in accordance with the ACMG-AMP guidelines. EVE, CV disagree: Total number of variants for which the EVE model predicts Benign/Pathogenic when the label in ClinVar is Pathogenic/Benign. EVE _and_ACMG, CV disagree: Total number of variants for which the EVE model, when combined with other evidence in accordance with the ACMG-AMP criteria, predicts Benign/Pathogenic when the label in ClinVar is Pathogenic/Benign. B_LB Pre: Number of (Likely) Benign labels with at least as one-star rating. P_LP Pre: Number of (Likely) Pathogenic labels with at least as one-star rating. U pre: Number of variants of uncertain significance, as obtained from ClinVar and gnomAD. (B, P, U) mod: Number of variants predicted Benign, Pathogenic or Uncertain respectively by the EVE model. (B_LB, P_LP, U) post: Number of variants predicted (Likely) Benign, (Likely) Pathogenic or Uncertain following our analysis combining EVE predictions with other sources of evidence in accordance with the ACMG-AMP guidelines.
